# GenoSim: A Forward-Time Genotype Simulator for Clinical and Population Genetics with Population Stratification

**DOI:** 10.64898/2026.06.20.733503

**Authors:** Abu Bakar, Rutaba Gul, Tayyab Afghani, Saad Serfraz

## Abstract

**Motivation:** Next-generation sequencing studies in clinical genetics are often limited by the scarcity of human genotype data, which stems from ethical, regulatory, and economic barriers. The shortfall is sharpest in consanguineous populations, which are common in South Asia and the Middle East, where family-based designs need large pedigrees that are rarely sequenced in full. Existing simulators do not combine pedigree-aware propagation, realistic population stratification, and clinical export formats in one tool.

**Results:** We present GenoSim, an R package for forward-time simulation of diploid SNP genotypes. It runs in two modes: a population mode implementing inbreeding-adjusted Hardy-Weinberg sampling, Wright-Fisher drift, directional selection, recurrent mutation, and Haldane recombination across multiple generations; and a pedigree-constrained mode that ingests real family VCFs and a pedigree, reconstructs phase where the pedigree makes it identifiable, propagates genotypes through the observed family structure, and appends synthetic generations. Version 1.1.1 adds population stratification through the Balding-Nichols model parameterised by gnomAD v3.1 fixation indices (*F_sr_*) for eight ancestry groups (AFR, AMR, EAS, EUR, FIN, MID, SAS, ASJ), empirical allele-frequency loading from external reference panels, and admixed-cohort simulation. Analysis functions cover Hardy-Weinberg testing, linkage disequilibrium, runs of homozygosity, principal component analysis, founder-referenced and between-generation *F*-statistics, and Nei gene diversity. Output is compatible with VCFv4.2, PLINK PED/MAP/RAW, and tidy CSV.

**Availability and implementation:** GenoSim is available as an R package at https://github.com/malikbak/GenoSim under the MIT licence. It requires R ≥4.0.0 and depends only on base R packages (stats, utils, graphics, grDevices, tools).

## 1 Introduction

The rapid decrease in sequencing costs has made whole-genome and whole-exome sequencing far more accessible, yet clinical genetics research in many settings remains severely data-limited. Ethical constraints on sharing patient genomes, small family cohort sizes, and the expense of large-scale population biobanks create a persistent bottleneck: researchers cannot validate computational pipelines, estimate statistical power, or train machine learning models without adequate training data [1,2]. This challenge is particularly acute in consanguineous populations, which are prevalent across South Asia, the Middle East, and North Africa, where autosomal recessive disorders arise at elevated rates and family-based autozygosity mapping is a primary gene discovery strategy[3].

Existing genotype simulators address different parts of this problem. HAPGEN2[4] and fastPHASE[5] simulate haplotypes from reference panels but do not model pedigree structures. PLINK[6] can generate synthetic genotypes under simplified population models, while forward-time simulators such as SLiM[7] and msprime[8] offer population-level realism but require considerable bioinformatics expertise. None provides integrated pedigree-constrained propagation from real observed VCFs together with ancestry-specific allele frequency priors in a single, clinical-ready R package.

Here we introduce GenoSim, an R package developed to address this gap. GenoSim lets researchers seed simulations from a small observed family cohort, propagate genotypes through the exact pedigree structure with realistic recombination and inbreeding, and generate arbitrarily large synthetic cohorts whose allele frequency spectrum matches a target ancestral population. The package includes the analysis, visualisation, and export functions needed to produce VCF-ready data compatible with standard GWAS and rare variant association pipelines.

## 2 Methods

### 2.1 Overview and notation

GenoSim simulates diploid biallelic single-nucleotide polymorphism (SNP) genotypes. Each genotype is stored as an allele dosage *g* ∈ {0,1,2}, the count of alternate alleles carried at a locus. *A* generation is a matrix *G*^(*t*)^ of size *n*_*t*_ x *L*, where *n*_*t*_ is the number of individuals in generation *t* and *L* is the number of loci. We write *p*_*l,t*_ for the alternate-allele frequency at locus *l* in generation *t* and *q*_*l,t*_ = 1 − *P*_*l,t*_*·* Missing calls are permitted, and every estimator below ignores them locus by locus.

Two entry points share the same forward machinery. Population mode starts from founder allele frequencies. Pedigree mode starts from observed family genotypes and a pedigree, reconstructs the observed generations, and then continues forward under the same model.

### 2.2 Founder genotypes

Founder alternate-allele frequencies are drawn independently per locus. The minor-allele frequency is sampled uniformly on [maf_min_, 0.5], and the alternate label is assigned to the minor or major allele with equal probability. Given a founder frequency *p* and an inbreeding coefficient *F* ∈ [0,1), each founder genotype is sampled from the inbreeding adjusted Hardy-Weinberg distribution

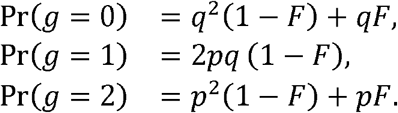

At *F* = 0 this reduces to Hardy-Weinberg proportions [9]. For *F* > 0 the heterozygote class is deflated by a factor 1 - *F* and the two homozygote classes absorb the difference, which is the standard model for a population mating at rate *F* among relatives [10].

### 2.3 SNP positions and the recombination map

Each locus is placed on an autosome. The number of loci assigned to a chromosome is proportional to its physical length, and positions are sampled without replacement along the chromosome, so a SNP identifier and its recorded base-pair coordinate always agree. For two adjacent markers separated by *d* base pairs on the same chromosome, the per meiosis recombination probability follows the Haldane mapping function [l l],

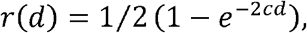

where *c* is the recombination rate per base pair (default *c* = 10^−8^). Markers on different chromosomes segregate independently.

### 2.4 Gamete transmission

*A* gamete is a haploid allele vector passed from one parent. Let the parent’s two haplotypes be *h*^(1)^ and *h*^(2)^. The gamete is built one chromosome at a time. The copy used at the first marker of a chromosome is chosen with probability 1/2 for each haplotype. Moving along the chromosome, the copy switches between the two haplotypes at marker *k* + 1 with probability *r(d*_*k,k+1*_) and otherwise stays on the current haplotype, which reproduces crossover events at the per-interval rate set by the Haldane map. Because phase is only partially known from unphased dosages, the haplotypes used for transmission are reconstructed *as* 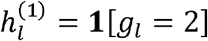 and 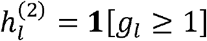. This preserves the dosage at every locus and fixes a within-parent phase; pedigree mode refines it wherever parental genotypes make phase identifiable (Section 2.8).

An offspring genotype is the sum of two independent gametes, one from each parent,

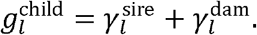

When the mating is inbred with offspring coefficient *F*_*i*_, the two gametes are made identical with probability *F*_*i*_, so the offspring receives two copies of the same transmitted haplotype. This produces the expected excess of homozygosity.

### 2.5 Drift, mutation, and selection

Drift arises from finite sampling rather than from perturbing allele frequencies directly. At each generation the breeding pool is capped at the effective size *N*_*e*_: when more individuals are available, *N*_*e*_ of them are drawn at random to be parents. Random mating among these breeders, followed by transmission, makes the allele count in the next generation a finite sample of the parental gene pool, so the expected loss of gene diversity per generation matches the Wright-Fisher rate 1/(2*N*_*e*_) [12](Section 2.9).

Recurrent two-allele mutation is applied to each transmitted gamete: at every locus the transmitted allele is flipped, 0↔1, independently with probability *µ* per allele per generation. This reintroduces variation at loci that have drifted towards fixation.

Selection acts on whole genotypes as viability selection, which keeps linkage between loci intact Each locus contributes an additive fitness factor under the model *w*_*AA*_ = l +*s, w*_*aa*_ = 1 + *s*/2, *w*_*aa*_ = l, where s is the selection coefficient on the alternate allele[13]. Multilocus fitness is the product over loci, computed on the log scale for numerical stability,

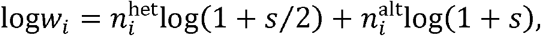

where 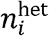 and 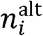 count the heterozygous and alternate-homozygous loci in individual *i*. Offspring then survive in proportion to relative fitness; the next generation is formed by sampling with replacement using probabilities

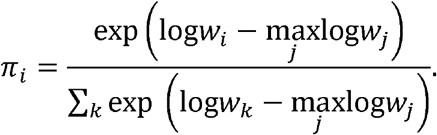

At *s* = 0 the weights are uniform and selection has no effect

### 2.6 Allele-frequency bookkeeping

The allele frequency recorded for a generation is the realised frequency of its stored genotypes,

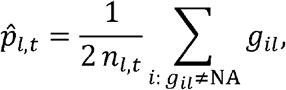

where *n*_*l* ,*t*_ is the number of non-missing calls at locus *l*. Drift, mutation, and selection all act on the genotypes themselves, so the frequency trajectory and every summary statistic stay consistent with the matrices returned to the user.

### 2.7 Population stratification

Structured populations use the Balding-Nichols model [14]. Given an ancestral frequency *p*_*A*_ and a population-specific fixation index *F*_*sT*_, the population frequency at a locus is drawn from a Beta distribution,

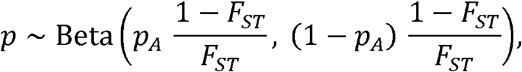

with mean E[*p*] = *p*_*A*_ and variance Var[*p*] = *p*_*A*_ (1−*p*_*A*_) *F*_*sT*_ *·* Genotypes are then sampled from the inbreeding-adjusted Hardy-Weinberg distribution of Section 2.2 using *p*. Built-in *F*_*sT*_ values for major continental groups are provided (Table **1)**.

### 2.8 Pedigree-constrained simulation

#### 2.8.1 Kinship and inbreeding coefficients

Inbreeding coefficients are obtained from the pedigree by the recursive tabular method [15]. Let *φ(X, Y)* be the coancestry (kinship) of individuals *X* and *Y*. With parents ordered before offspring, for a non-founder *X* with parents *P*_*x*_ and *M*_*x*_,

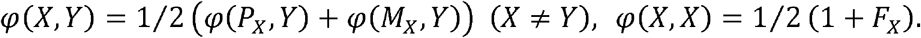

The inbreeding coefficient of an individual is the coancestry of its parents,

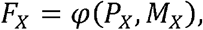

with founders set to *F* = 0 and *φ (X, X)* = 1/2. The offspring of first cousins has *F* = 1/16, and the offspring of a double-first-cousin or uncle-niece union has *F* = 1/8.

#### 2.8.2 Pedigree-informed phasing

A genotyped individual with both parents genotyped is phased from the transmission implied by the parents. Where the child is heterozygous and one parent is homozygous, parental origin is fixed: the homozygous parent can transmit only its single allele type, so the other allele must come from the second parent. When both parents are heterozygous, or a parent is missing, the phase at that locus is left random. Phase is therefore resolved exactly at the informative sites and falls back to a coin flip only where the data are uninformative.

#### 2.8.3 Reconstruction of observed generations

Observed generations are filled in pedigree order. A genotyped individual keeps its observed dosage vector. A non-genotyped individual with both parents available is generated by the transmission step above, with its pedigree *F*_*i*_ controlling the selfing probability and mutation applied to its gametes. A non-genotyped individual without usable parents is imputed by sampling from the inbreeding-adjusted Hardy-Weinberg distribution at the relevant allele frequencies, taken in order of preference from a supplied population-stratified set, an external reference, or the current cohort. When imputation of a read-in cohort is requested, any residual missing calls are filled from the per-locus allele frequency, with a global-frequency fallback for loci that have no observed call, so the returned matrix is complete.

#### 2.8.4 Synthetic descendant generations

Once the observed generations are reconstructed, further generations are appended under the same forward model: finite-pool random mating capped at *N*_*e*_, recombinant transmission, mutation, and viability selection. The allele frequency recorded for each synthetic generation is again the realised frequency of its genotypes.

### 2.9 Heterozygosity and inbreeding statistics

#### 2.9.1 Observed and expected heterozygosity

The observed heterozygosity of generation *t* is the fraction of heterozygous calls,

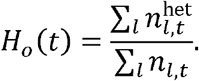

Expected heterozygosity (Nei gene diversity) is estimated per locus with the finite-sample correction, which matters for the small samples typical of a family [16],

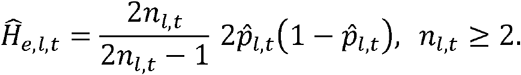

Using the uncorrected 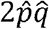 biases 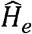 downwards and, in small samples, can push the index of the next subsection below zero by roughly − (1−*F*_*IS*_)/(2*n* − 1).

#### 2.9.2 Within-generation inbreeding

The within-generation index is Wright’s *F*_*IS*_ referenced to the current generation’s own frequencies,

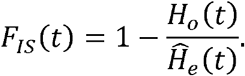

This measures departure from Hardy-Weinberg proportions within generation *t*. A single round of random mating restores those proportions, so *F*_*IS*_ returns towards zero after inbred founders mate, however inbred they were. It does not accumulate and is not a measure of cumulative inbreeding.

#### 2.9.3 Founder-referenced inbreeding

To capture inbreeding that builds up over generations, the founder gene pool is used as a fixed baseline. With 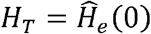 in the founders, and summing over loci polymorphic in the founders (a ratio of sums, which stays stable at low-frequency loci),

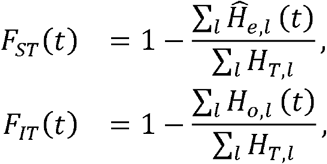

with the within-generation index of the previous subsection completing the Wright identity,

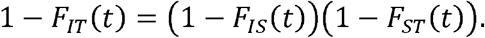

Because *F*_*ST*_(*t*) and *F*_*IT*_(*t*) are measured against the fixed founders, they accumulate as drift erodes diversity rather than returning to zero: *F*_*ST*_(*t*) is the cumulative loss of gene diversity since the founders, and *F*_*IT*_(*t*) is the total inbreeding of current individuals relative to the founder gene pool.

#### 2.9.4 Expected drift trajectory and effective size

Under Wright-Fisher drift in a population of effective size *Ne*, two gene copies coalesce in the previous generation with probability 1/(2*N*_*e*_), which gives the gene-diversity recurrence

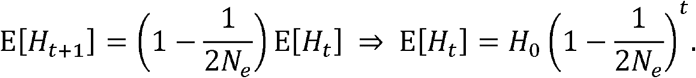

The complementary inbreeding recurrence gives the expected founder-referenced curve, with *F*_0_ = 0,

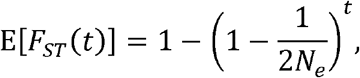

which GenoSim reports next to the simulated *F*_*ST*_(*t*) as a check. The realised effective size is estimated from the diversity decay between consecutive generations,

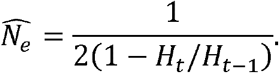

The mean pedigree inbreeding of a generation, *F*_ped_(*t*) =1/*n*_t_ Σ*i*∈*t Fi*, is reported as a frequency-free comparison and predicts E[*H*_*0*_(*t*)] = (1 – *F*_ped_ (*t*)*) H*_*T*_,

### 2.10 Downstream analyses

#### 2.10.1 Hardy-Weinberg equilibrium test

Each locus is tested with a one-degree-of-freedom chi-squared goodness-of-fit statistic. Let *n*_*0*_, *n*_*1*_,*n*_*2*_ be the observed genotype counts and *n*_*c*_ = *n*_*0*_ + *n*_1_ + *n*_2_ the number of called genotypes. With 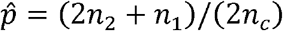 and 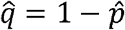, the expected counts are 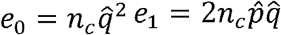, and 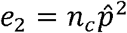, and

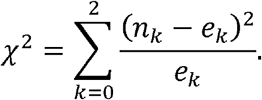

Using the called count *n*_*c*_ rather than the total number of individuals keeps the observed and expected totals equal, so missing data do not inflate the statistic. Fixed loci are reported as such rather than tested.

#### 2.10.2 Linkage disequilibrium

For a pair of loci with dosage vectors *x* and *y* over the individuals genotyped at both, *r*^2^ is the squared Pearson correlation of the dosages, the standard genotypic estimate [17],

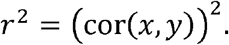

The haplotypic coefficient *D* is recovered from the dosage covariance through cov (*x, y*)= 2D|,so 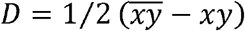. With allele frequencies *p*_*x*_ = *x/2* and *p*_*y*_*= y*/2, the normalised coefficient is (Lewontin, 1964)

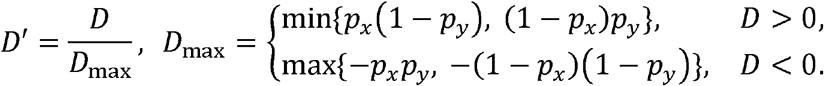

On this scale two perfectly correlated SNPs give *r*^*2*^ = *D*′ = 1. Pairs are evaluated within a chromosome up to a maximum separation, and monomorphic loci are skipped.

#### 2.10.3 Runs of homozygosity

For each individual, every chromosome is scanned for contiguous runs of homozygous calls. A small number of heterozygous or missing calls is tolerated inside a run before it is broken. A run is reported when it spans at least a minimum number of SNPs and a minimum physical length. The per-individual run burden, and the genomic inbreeding coefficient *F*_RoH_ derived from the fraction of the autosome that lies in long runs, are the standard way to detect recent common ancestry, because recent identity by descent leaves long runs while the homozygosity from background drift is short and scattered[18].

#### 2.10.4 Principal component analysis

Genomic structure is summarised by PCA on the dosage matrix after loci with zero variance are removed [19]. Component scores, loadings, and the percentage of variance explained, 100 *‘λ*_*k*_**/***Σ*_*j*_*λ*_*j*_with*‘λ*_*k*_ the *k-th* eigenvalue, are returned for the requested number of components.

#### 2.10.5 Between-generation differentiation

Differentiation between two generations is estimated with a Weir-Cockerham formulation[20]. For sample sizes *n*_1_, *n*_*2*_. and allele frequencies *p*_*1*_, *p*_*2*_ at a locus, with pooled frequency *p* = (*n*_1_*p*_1_ + *n*_2_*p*_2_)/(*n*_1_ + *n*_2_),

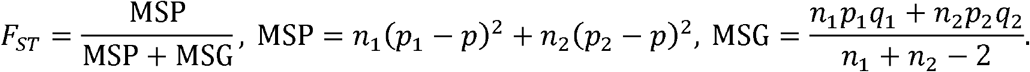

The mean and median across loci are reported for each pair of generations.

#### 2.10.6 Diversity statistics

Per generation, Nei gene diversity is 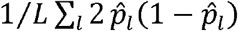, the proportion of segregating sites is the fraction of loci with 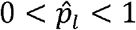, and Watterson’s estimator is 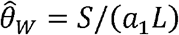 with *S* the number of segregating sites and 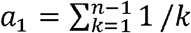.

### 2.11 Implementation

Geno Sim is written in R and depends only on the base distribution. All sampling steps are seedable, so a run reproduces exactly from its seed. Genotype matrices may contain missing values, and every estimator above is computed on the non-missing entries. For inputs too large to hold in memory, a streaming pre-filter reduces a VCF to a biallelic autosomal subset before the in-memory reader runs.

## 3 Results and Discussion

### 3.1 Study family and simulation setup

We applied GenoSim in pedigree mode to a real family with cousin marriages, using the whole VCF with no allele-frequency filter. GenoSim retained 24,024 biallelic autosomal SNPs. Generations 0 to 4 are the observed pedigree, with 7, 2, 8, 18, and 7 genotyped individuals, and generations 5 to 13 are synthetic descendant generations of 9 individuals each. The drift benchmark uses Ne = 7, since the expected founder-referenced FST at generation 1 is 0.07143 = 1/(2 × 7). Tables 2 and 3 give the per-generation summaries and the founder-referenced inbreeding statistics, and Figures 1 and 2 show the heterozygosity and inbreeding trajectories.

**Table 1.** gnom AD v3.1 population parameters used in GenoSim.

| Populatio<br>n | Code | $F_{ST}$ | Mean Het | gnomAD | |
| --- | --- | --- | --- | --- | --- |
| | | | | $N$ | Mean MAF |
| African/Af<br>r.-<br>American | AFR | 0.155 | 0.352 | 20,744 | 0.198 |
| Admixed<br>American | AMR | 0.068 | 0.301 | 7,647 | 0.156 |
| East Asian | EAS | 0.108 | 0.272 | 9,977 | 0.138 |
| European<br>(non-Finn.) | EUR | 0.064 | 0.286 | 34,029 | 0.148 |
| Finnish | FIN | 0.091 | 0.258 | 5,316 | 0.131 |
| Middle<br>Eastern | MID | 0.072 | 0.291 | 158 | 0.152 |
| South<br>Asian | SAS | 0.082 | 0.279 | 15,308 | 0.144 |
| Ashkenazi | ASJ | 0.085 | 0.261 | 5,076 | 0.134 |

| Populatio |  |  |  | gnomAD |  |
| --- | --- | --- | --- | --- | --- |
| n | Code | $F_{ST}$ | Mean Het | $N$ | Mean MAF |
| Jewish |  |  |  |  |  |

**Table 2.** Per-generation genotype summaries. Every generation has 24,024 SNPs.

| Gen | Source | n | Ho | He | FIS | Mean MAF | Frac. fixed |
| --- | --- | --- | --- | --- | --- | --- | --- |
| 0 | observed | 7 | 0.2179 | 0.1984 | -0.09826 | 0.1684 | 0.5701 |
|  |  |  | 3 | 3 |  | 8 | 0 |
| 1 | observed | 2 | 0.1983<br>4 | 0.1491<br>4 | -0.32989 | 0.1225<br>2 | 0.6484<br>8 |
| 2 | observed | 8 | 0.1831<br>6 | 0.1860<br>7 | 0.01566 | 0.1503<br>5 | 0.5708<br>5 |
| 3 | observed | 18 | 0.2172<br>0 | 0.2056<br>6 | -0.05611 | 0.1798<br>6 | 0.5694<br>7 |
| 4 | observed | 7 | 0.2170<br>6 | 0.1983<br>2 | -0.09451 | 0.1681<br>9 | 0.5701<br>8 |
| 5 | synthetic | 9 | 0.2121<br>3 | 0.1892<br>6 | -0.12083 | 0.1553<br>7 | 0.5699<br>7 |
| 6 | synthetic | 9 | 0.2095<br>6 | 0.1788<br>0 | -0.17207 | 0.1438<br>3 | 0.5756<br>7 |
| 7 | synthetic | 9 | 0.1969<br>8 | 0.1666<br>7 | -0.18185 | 0.1313<br>8 | 0.5855<br>4 |
| 8 | synthetic | 9 | 0.1549<br>2 | 0.1592<br>0 | 0.02688 | 0.1242<br>0 | 0.5969<br>4 |
| 9 | synthetic | 9 | 0.1782<br>8 | 0.1459<br>5 | -0.22157 | 0.1134<br>3 | 0.6188<br>0 |
| 10 | synthetic | 9 | 0.1516<br>8 | 0.1404<br>3 | -0.08009 | 0.1087<br>7 | 0.6316<br>6 |
| 11 | synthetic | 9 | 0.1600<br>7 | 0.1296<br>7 | -0.23438 | 0.1008<br>4 | 0.6593<br>4 |
| 12 | synthetic | 9 | 0.1290<br>1 | 0.1212<br>5 | -0.06401 | 0.0935<br>2 | 0.6794<br>9 |
| 13 | synthetic | 9 | 0.1298<br>2 | 0.1125<br>9 | -0.15308 | 0.0857<br>6 | 0.6952<br>6 |

**Table 3.**
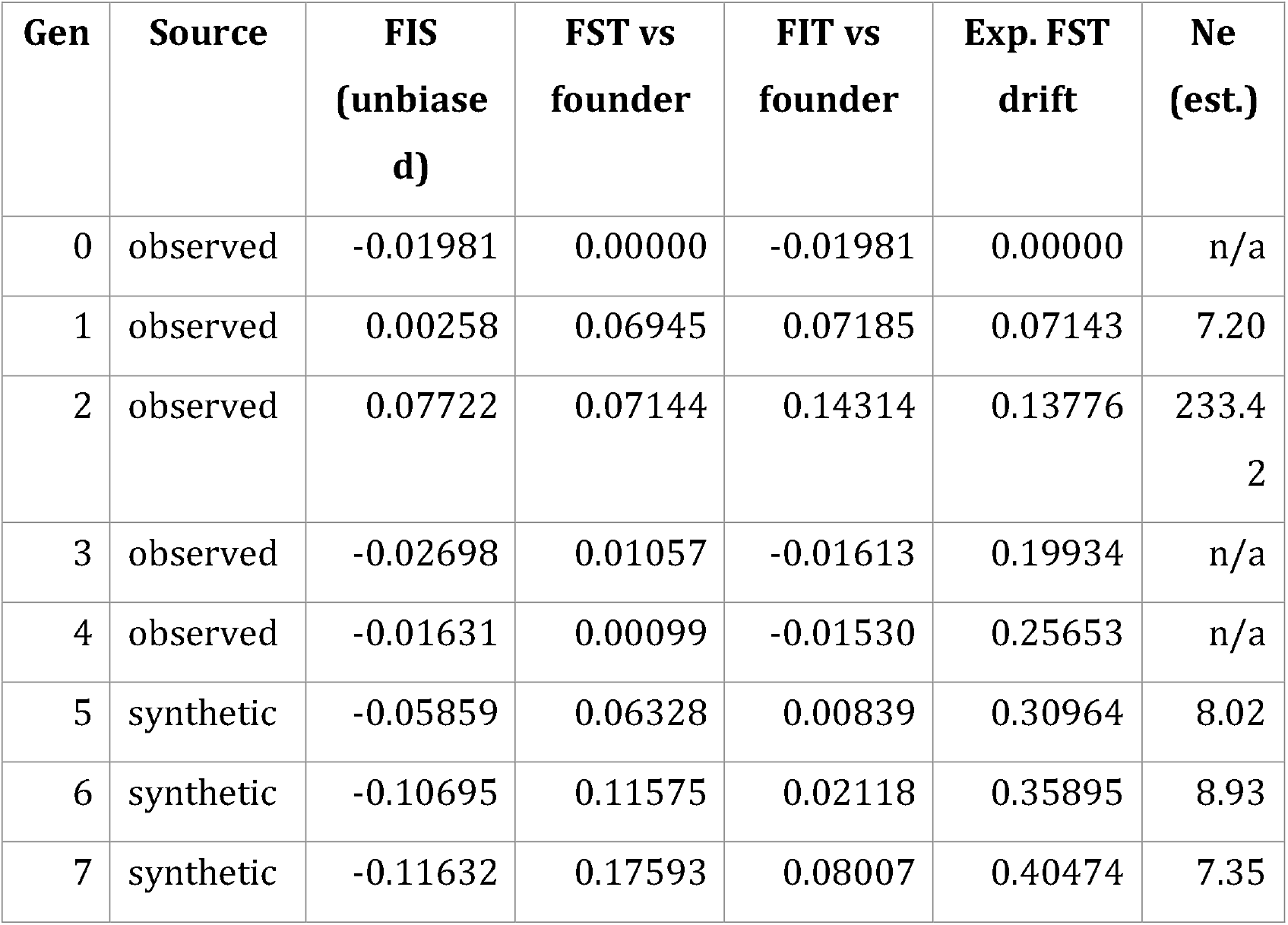
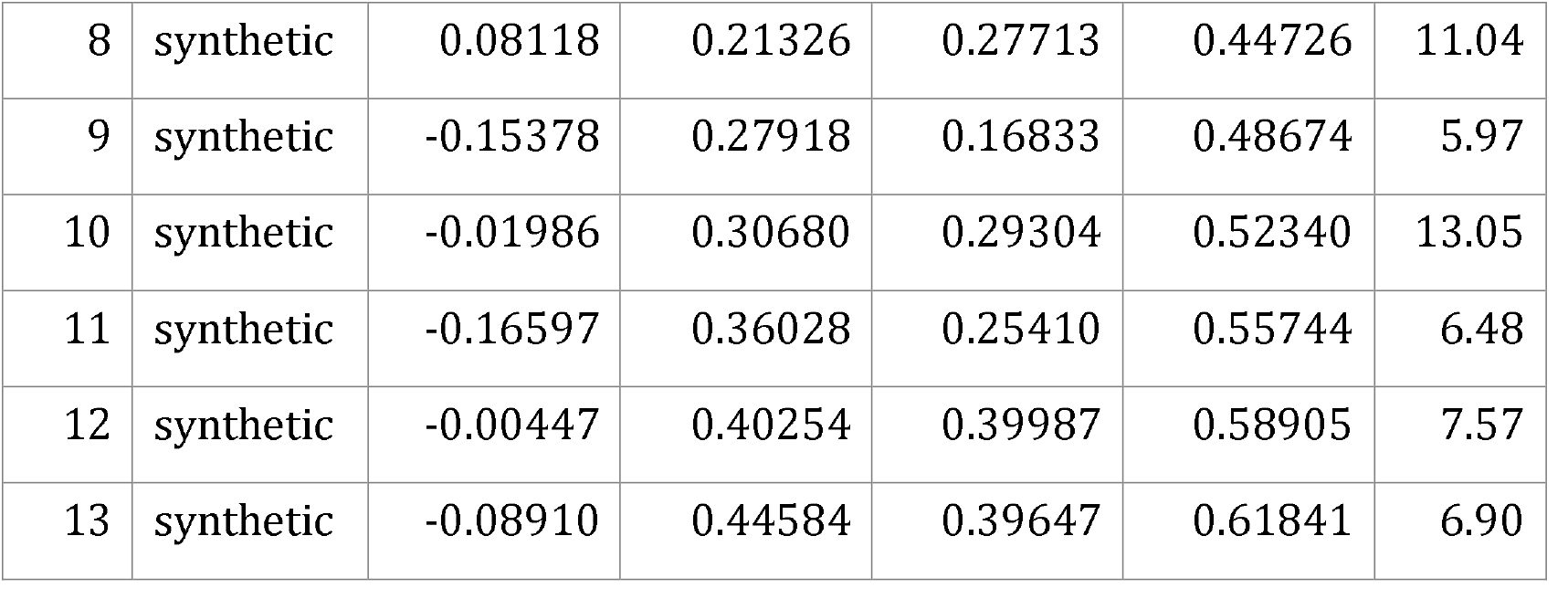
Founder-referenced inbreeding statistics. “n/a” marks an undefined value.

| Gen | Source | FIS<br>(unbiased) | $F_{ST}$ vs<br>founder | $F_{IT}$ vs<br>founder | Exp. $F_{ST}$<br>drift | $N_e$<br>(est.) |
| --- | --- | --- | --- | --- | --- | --- |
| 0 | observed | -0.01981 | 0.00000 | -0.01981 | 0.00000 | n/a |
| 1 | observed | 0.00258 | 0.06945 | 0.07185 | 0.07143 | 7.20 |
| 2 | observed | 0.07722 | 0.07144 | 0.14314 | 0.13776 | 233.4<br>2 |
| 3 | observed | -0.02698 | 0.01057 | -0.01613 | 0.19934 | n/a |
| 4 | observed | -0.01631 | 0.00099 | -0.01530 | 0.25653 | n/a |
| 5 | synthetic | -0.05859 | 0.06328 | 0.00839 | 0.30964 | 8.02 |
| 6 | synthetic | -0.10695 | 0.11575 | 0.02118 | 0.35895 | 8.93 |
| 7 | synthetic | -0.11632 | 0.17593 | 0.08007 | 0.40474 | 7.35 |
| 8 | synthetic | 0.08118 | 0.21326 | 0.27713 | 0.44726 | 11.04 |
| 9 | synthetic | -0.15378 | 0.27918 | 0.16833 | 0.48674 | 5.97 |
| 10 | synthetic | -0.01986 | 0.30680 | 0.29304 | 0.52340 | 13.05 |
| 11 | synthetic | -0.16597 | 0.36028 | 0.25410 | 0.55744 | 6.48 |
| 12 | synthetic | -0.00447 | 0.40254 | 0.39987 | 0.58905 | 7.57 |
| 13 | synthetic | -0.08910 | 0.44584 | 0.39647 | 0.61841 | 6.90 |

**Figure 1.**
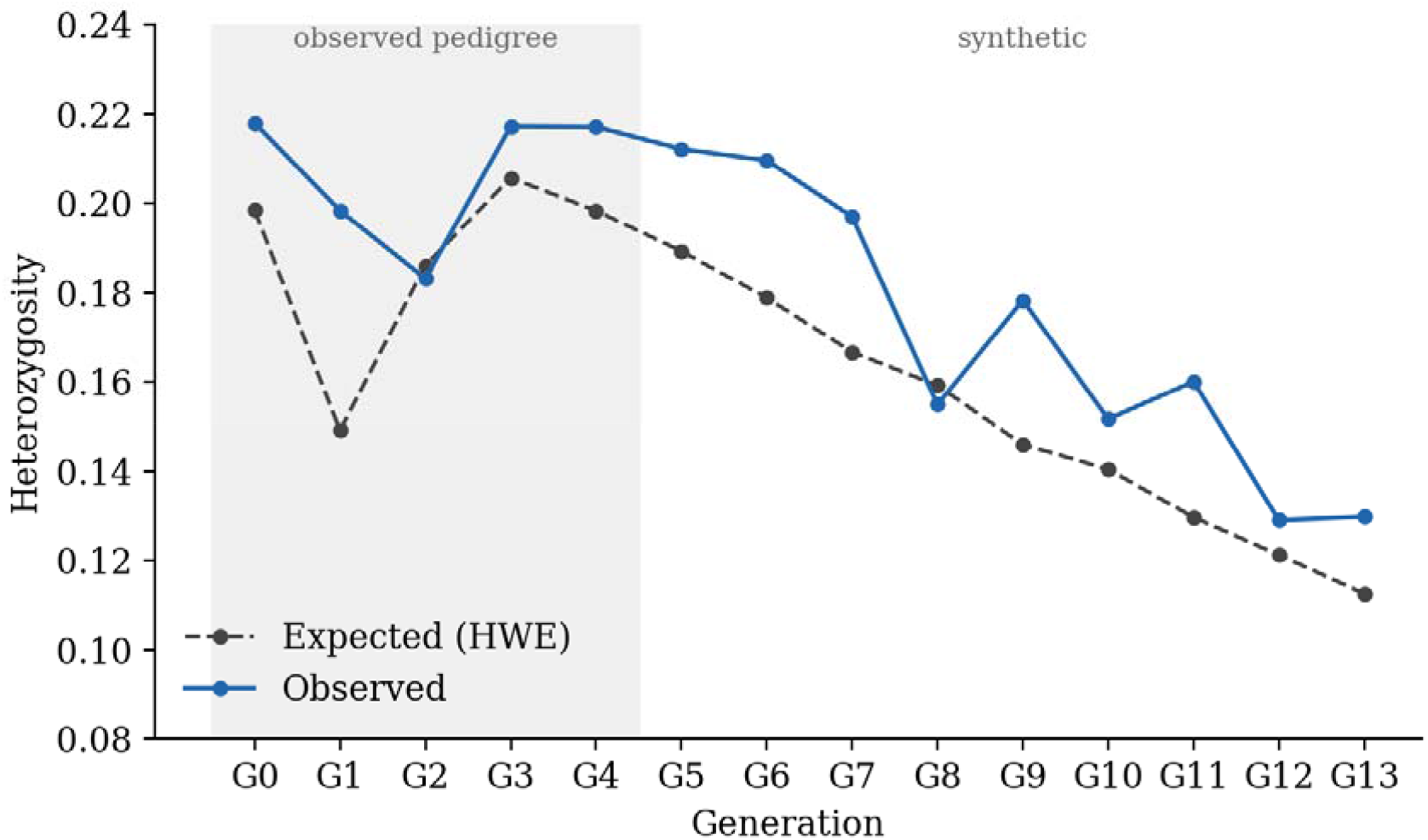
Observed and expected (Hardy-Weinberg) heterozygosity across generations. Shaded region marks the observed pedigree generations; later generations are synthetic. Per generation FIS values are given in Table 2.

**Figure 2.**
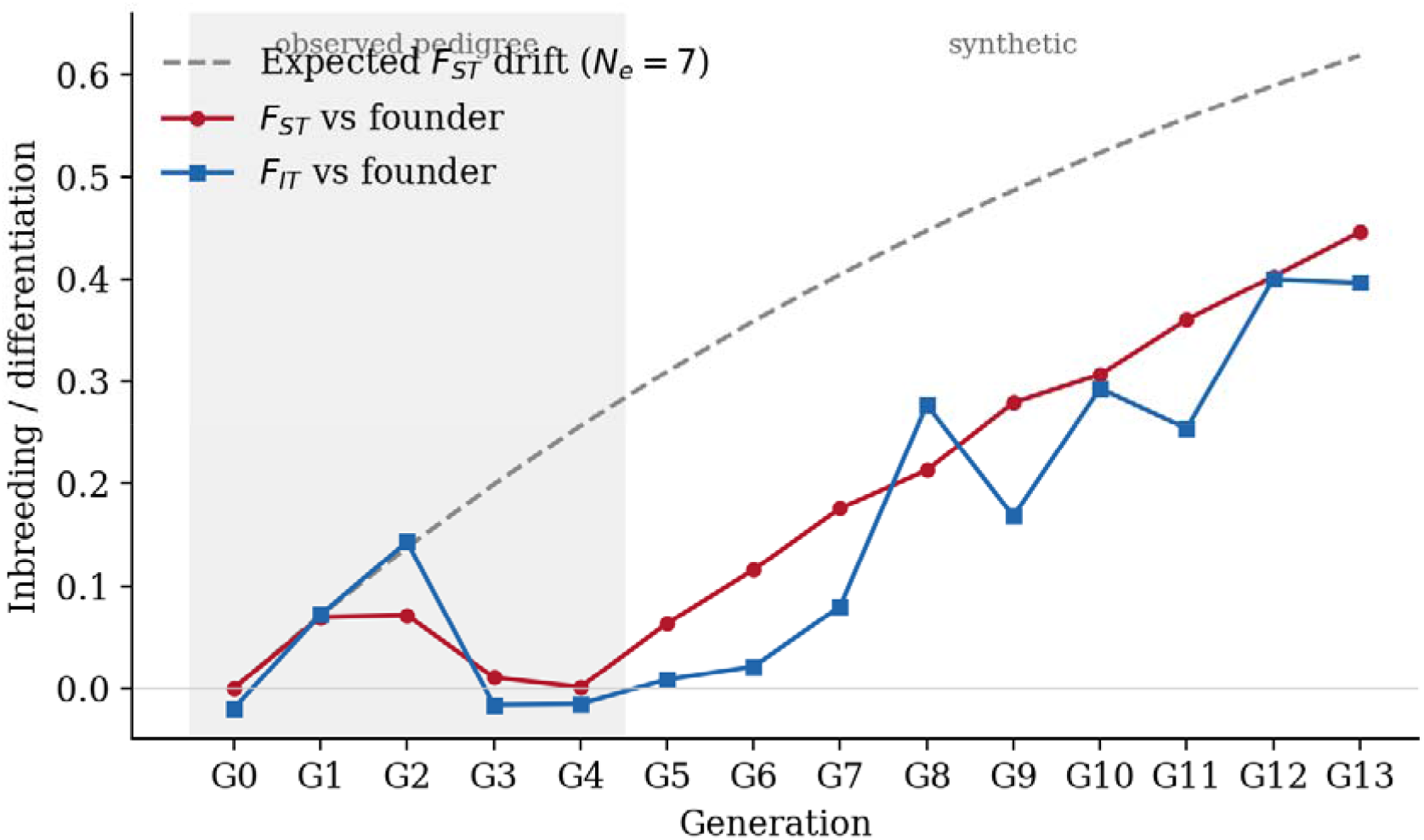
Founder-referenced FST and FIT against the expected Wright-Fisher drift curve for Ne= 7. Shaded region marks the observed pedigree generations.

### 3.2 Heterozygosity, founder-referenced inbreeding, and drift

Heterozygosity falls steadily across the pedigree (Figure 1, Table 2). Observed heterozygosity declines from 0.218 in the founders to 0.130 at generation 13, about 40%, and expected heterozygosity from 0.198 to 0.113, about 43%. Over the same span the mean minor-allele frequency falls from 0.168 to 0.086 and the fraction of fixed sites rises from 0.57 to 0.70. Nei gene diversity tracks expected heterozygosity, the fraction of segregating sites falls from 0.43 to 0.30, and Watterson’s theta falls from 0.174 to 0.112. The lineage loses roughly two fifths of its founding diversity over fourteen generations.

The within-generation inbreeding coefficient FIS scatters near zero with no trend (Table 2), ranging from -0.33 to 0.03 in the raw estimate and behaving the same way after bias correction (Table 3). This is expected rather than contradictory. FIS is referenced to each generation’s own allele frequencies, so a single round of random mating restores Hardy Weinberg proportions and the index resets toward zero; it cannot accumulate. The observed pedigree generations are also small, 2 to 18 individuals, where FIS is noisy. The cumulative signal of inbreeding lives instead in the founder-referenced statistics.

Measured against the fixed founder gene pool, inbreeding accumulates as expected (Figure 2, Table 3). The founder-referenced FST rises almost monotonically to 0.446 by generation 13 and FIT to 0.396. Both sit below the expected Wright-Fisher drift curve for Ne= 7, which reaches 0.618 at generation 13, because the realised effective size is a little larger than the benchmark. The heterozygosity decay across the synthetic phase implies a per-generation retention of about 0.937, that is 1 - 1/(2 Ne) with Ne close to 8. At that effective size the inbreeding added each generation is about 1/(2 Ne), or 1/16, which is 0.063: the inbreeding coefficient of a first-cousin child. Compounded across the nine synthetic generations this reaches a cumulative value near 0.44, in the range of the reported FST and FIT. Between consecutive generations the Weir-Cockerham FST averages 0.073, and the founder-to-final differentiation is 0.24 on the same scale.

### 3.3 Runs of homozygosity confirm recent identity by descent

The genome-wide statistics above show that the lineage becomes more homozygous, but they cannot on their own show that the homozygosity is recent identity by descent rather than background drift. Runs of homozygosity settle that question, because recent common ancestry leaves long homozygous segments while drift leaves short scattered ones. In the observed generations the mean genomic inbreeding coefficient FROH ranges from 0.074 to 0.095 (Table 4), and per-individual values span 0.063 to 0.134. These are the values expected for the offspring of close relatives: a first-cousin child carries about 0.063, and the most homozygous individual in the pedigree, in generation 2, reaches 0.134, near the level of a double-first-cousin or uncle-niece union. The FROH burden is highest in generation 2, consistent with the closest consanguinity falling in that part of the pedigree.

**Table 4.** Genomic inbreeding coefficient FROH by generation, computed on the observed pedigree individuals.

| Gen | Mean $F_{ROH}$ | n individuals |
| --- | --- | --- |
| 0 | 0.0736 | 7 |
| 2 | 0.0948 | 4 |
| 3 | 0.0773 | 9 |
| 4 | 0.0811 | 5 |

### 3.4 Reading the same signatures two ways

The same numbers admit two readings that the genome-wide summaries cannot separate. Read as a consanguineous family, the trajectory is sustained inbreeding at roughly cousin level per generation: heterozygosity falls, fixed sites accumulate, and the founder referenced FST and FIT climb to about 0.4, so each generation carries less of the founders’ diversity and more homozygosity. Read as a general population, the identical numbers are ordinary Wright-Fisher drift in a small isolate: expected heterozygosity decays geometrically, FST accumulates as temporal differentiation from the founder pool, and alleles drift to fixation because only about eight individuals breed each generation. Consanguinity is inbreeding at the level of specific matings; small-population drift is inbreeding at the level of the whole gene pool. Both lower heterozygosity and both raise FST and FIT.

Two further signatures are consistent with both readings but help anchor the interpretation. Departures from Hardy-Weinberg proportions in the observed generations are modest and non-monotonic, with 4 to 13 percent of tested sites failing the test across generations 0 to 4, in line with the near-zero FIS and the small sample sizes rather than a systematic heterozygote deficit. Linkage disequilibrium, by contrast, extends over long distances: several SNP pairs separated by more than half a megabase remain in near complete association, with D-prime close to 1 and r-squared above 0.8 (Table 5). Extended long-range LD of this kind is the expected footprint of recent identity by descent, since the shared haplotype blocks introduced by cousin marriage have had few meioses in which to recombine.

**Table 5.** Representative long-range linkage-disequilibrium pairs, showing near-complete association across large physical distances.

| Chr | SNP A<br>(Mb) | SNP B<br>(Mb) | Distance<br>(kb) | $r^2$ | $D'$ |
| --- | --- | --- | --- | --- | --- |
| 19 | 40.32 | 40.88 | 554 | 0.833 | 1.000 |
| 11 | 2.13 | 1.24 | 889 | 0.804 | 1.000 |
| 13 | 98.45 | 98.80 | 344 | 0.804 | 1.000 |
| 11 | 14.27 | 14.83 | 567 | 0.583 | 1.000 |
| 17 | 77.50 | 78.36 | 858 | 0.583 | 1.000 |

The feature that separates the two readings is the length of the homozygous segments. A runs-of-homozygosity analysis, not the genome-wide averages, is what decides whether the homozygosity in a real sample comes from recent cousin ancestry or from a small ancestral population. For clinical genetics the relevant consequence is autosomal recessive disease: a child of related parents can be homozygous for a rare deleterious variant inherited from a common ancestor, and the risk grows with the inbreeding coefficient. The unfiltered marker set used here is the right input for that question, because an allele-frequency filter would remove the rare variants that drive recessive disease. The forward synthetic generations are best read as a scenario for how homozygosity grows if consanguineous mating continues, not as a risk estimate for any individual.

### 3.5 Comparison with existing tools

Table 6 sets GenoSim against related simulators. The combination it offers, pedigree constrained propagation from observed VCFs together with ancestry-specific founder priors in a dependency-free R package, is not available in any single existing tool.

**Table 6.** Feature comparison of GenoSim with related genotype simulation tools.

| Feature | GenoSim | SLiM | HAPGEN2 | PLINK | msprime |
| --- | --- | --- | --- | --- | --- |
| Pedigree input | Yes | No | No | Limited | No |
| VCF input | Yes | No | Yes | Yes | No |
| Population stratification | Yes (8 pops) | Yes | Partial | No | Yes |
| Inbreeding ($F$ model) | Yes | Yes | No | Limited | No |
| ROH / LD / PCA output | Yes | No | No | Yes | No |
| VCF / PLINK export | Yes | Yes | Yes | Yes | Yes |
| R package (no deps) | Yes | No | No | No | No |
| Clinical VCF FORMAT | Yes (DP/GQ/AD) | No | No | No | No |

## 4 Conclusion

GenoSim provides a unified, dependency-free R framework for generating realistic synthetic genotype data in clinical and population genetics. Its pedigree-constrained mode is built for studies where only a small family cohort is sequenced, and the Balding-Nichols stratification module makes simulated allele frequencies match the ancestry background of the study population. The worked family above shows how the same genome-wide summaries support a consanguinity reading and a small-population drift reading at once, with runs of homozygosity as the analysis that separates them. Together these features address the data scarcity that constrains rare disease genetics in under-represented populations. Planned extensions include Repp acceleration, phenotype simulation under disease models, pedigree-informed haplotype phasing, and integration with standard imputation and GWAS pipelines.

